# Hypoxia-induced mitoROS triggers M1 linear ubiquitin chains and activates NF-κB signaling

**DOI:** 10.1101/2025.11.06.686916

**Authors:** Kiri Tal, Jonathan Ram, Reis Noa, Maniv Inbal, Ordureau Alban, Michael H Glickman, Sulkshane Prasad

**Author notes:** Correspondence to Alban Ordureau, Prasad Sulkshane or Michael H Glickman. These authors contributed equally (Kiri Tal, Sulkshane Prasad).

## Abstract

Mitochondria are essential organelles responsible for cellular energy production and metabolism. Hypoxia, a pathophysiological condition, impairs the electron transport chain, disrupts mitochondrial function, and produces harmful reactive oxygen species (ROS). Ubiquitin signaling regulates mitochondrial health through several mechanisms, including protein degradation and mitophagy. Here, we show that hypoxia-induced mitophagy occurs independently of ubiquitination. However, mitochondria are heavily ubiquitinated under hypoxic stress. A significant portion of these hypoxia-induced ubiquitin chains constitute a specific type: linear head-to-tail fusions (M1), which are known for their role in NF-κB activation during cytokine signaling. We demonstrate that hypoxia-induced mitochondrial ROS leads to the accumulation of these M1 chains, activating NF-κB signaling and increasing the expression of its target genes. These findings reveal a critical internal signal that helps cells adapt to mitochondrial stress and triggers an inflammatory response.

## Introduction

Mitochondria are essential organelles responsible for harnessing molecular oxygen for cellular energy production and metabolism, signaling, and cell fate^1^. They function as a metabolic hub not only for oxidative phosphorylation but also for β-oxidation and the Krebs cycle, regulate intracellular calcium stores, determine cell fate through regulation of apoptosis, and regulate cellular redox homeostasis. Mitochondria are constantly exposed to various stresses, including unfolded protein response due to the complex nature of protein import from the cytosol, and to oxidative damage caused by the unavoidable production of ROS during oxidative phosphorylation^2^. Excess ROS build-up can cause oxidative damage to mitochondria, leading to their dysfunction^3^. When mitochondrial quality control fails, dysfunctional mitochondria accumulate and contribute to cellular damage, which is involved in many conditions like tumorigenesis, neurodegenerative diseases, stroke, and cardiomyopathies ^4^. Therefore, it is vital for the cell to either repair mitochondrial damage locally or remove severely damaged mitochondria altogether.

A mammalian cell has various quality control systems to address mitochondrial damage. Internal mitochondrial quality control via chaperones and proteases selectively refolds or degrades damaged mitochondrial proteins when the damage is localized^5^. When proteins in the mitochondrial outer membrane (MOM) are damaged, they are selectively degraded by the cytosolic ubiquitin proteasome system (UPS)^6,7^. The UPS also monitors mitochondrial protein import by modifying import substrates through the actions of E3 ubiquitin ligases and deubiquitinases (DUBs)^8,9^. This way, the UPS helps clear mislocalized intramitochondrial proteins and restores mitochondrial protein import by removing clogged import channels^9^. Irreversibly damaged mitochondria are ultimately eliminated by mitophagy^10^. Mitophagy is a specific autophagy process that targets damaged or unnecessary mitochondria for degradation to cellular homeostasis^11^. It can be mediated by either ubiquitin-dependent mechanisms (PINK1-Parkin mediated) or receptor-mediated ubiquitin-independent mechanisms^12^.

Attachment of ubiquitin to a target protein leads to various outcomes depending on the number of ubiquitin units, their orientation, and specific receptors for each lineage type. Ubiquitin can itself be ubiquitinated on one of the seven lysine (Lys) residues or at the N-terminus (M1), forming polyubiquitin chains that can encompass complex topologies^13^. Some chains are formed from repetitive homotypic attachments (single linkage type), while others are more complex and consist of heterogeneous (branched/mixed) ubiquitin chains^14^. Each linkage type encodes a different signaling message; for example, K48 typically targets proteins for proteasomal degradation, K63 is involved in vesicular trafficking, K6 has been linked to the PINK1-Parkin pathway for mitophagy, and Linear chains (M1) signal for NF-κB activation^14,15^. When located at mitochondria, each linkage type helps mitochondria respond differently to various stresses or metabolic changes.

Hypoxia hampers the mitochondrial respiratory chain due to decreased oxygen (O_2_) availability, the final electron acceptor. This results in inefficient electron transfer and the production of ROS by complexes I and III^16,17^. The generated ROS then damage macromolecules and other cellular structures, including the mitochondria themselves, rendering them dysfunctional^17^. We and others have shown that hypoxia-induced mitophagy occurs independently of ubiquitin receptors, mainly through specific MOM proteins like BNIP3 and NIX, which contain an LC-3-interacting region (LIR). These proteins facilitate the selective engulfment of damaged mitochondria by autophagosomes^18–20^. In parallel, mitochondria are also robustly ubiquitinated in response to hypoxia^20^.

In the present article, we further explore the importance of hypoxia-induced mitochondrial ubiquitination. We demonstrate that hypoxia-triggered mitochondrial ROS is the driving force for a number of parallel phenomena: HIF-1α stabilization, increased expression of BNIP3/NIX, induction of mitophagy, and mitochondrial ubiquitination, specifically M1 linear chains. Mitoquinone, a mitochondria-targeted antioxidant, blocked mitophagy and mitochondrial ubiquitination, indicating a complex relationship between mitochondrial ROS and ubiquitination. Treatment of hypoxic cells with either this mitochondrial antioxidant or Hoipin-8, an inhibitor for M1 chain assembly, depletes M1 Ub chains and inhibits NF-κB activation. Our study, therefore, reveals a role of mitochondria-derived ROS in activating an NF-κB signaling pathway and initiating an inflammatory response to hypoxic stress.

## Results

### Mitochondrial ubiquitination in response to hypoxia is unrelated to mitophagy

To evaluate the role of ubiquitination in hypoxia-triggered mitophagy, we monitored mitochondrial dynamics after exposure to hypoxia (24 hours at 1% oxygen). General autophagy was induced, as shown by the LC-3BI conversion to LC-3BII (**EV1A, EV1B**). Some LC-3BII-containing vesicles co-localized with fragmented mitochondria, suggesting mitochondrial-targeted autophagic vesicles (Mitophagy) **(EV1B)**. The extent of hypoxia-induced mitophagy was quantified using a mitochondrial matrix-targeted version of mtx-mKeima^XL^(^21^), referred to herein as mitoKeima, which shifts its fluorescence excitation wavelength when exposed to the acidic pH of lysosomes^22^. As anticipated, mitochondria became fragmented, and mitophagy increased in response to hypoxia **(Figure 1A, B)**. Additionally, immunoblotting analysis of isolated crude mitochondrial fractions showed significant accumulation of ubiquitin at 18 and 24 hours of hypoxia exposure **(Figure 1C)**.

**Figure 1:**
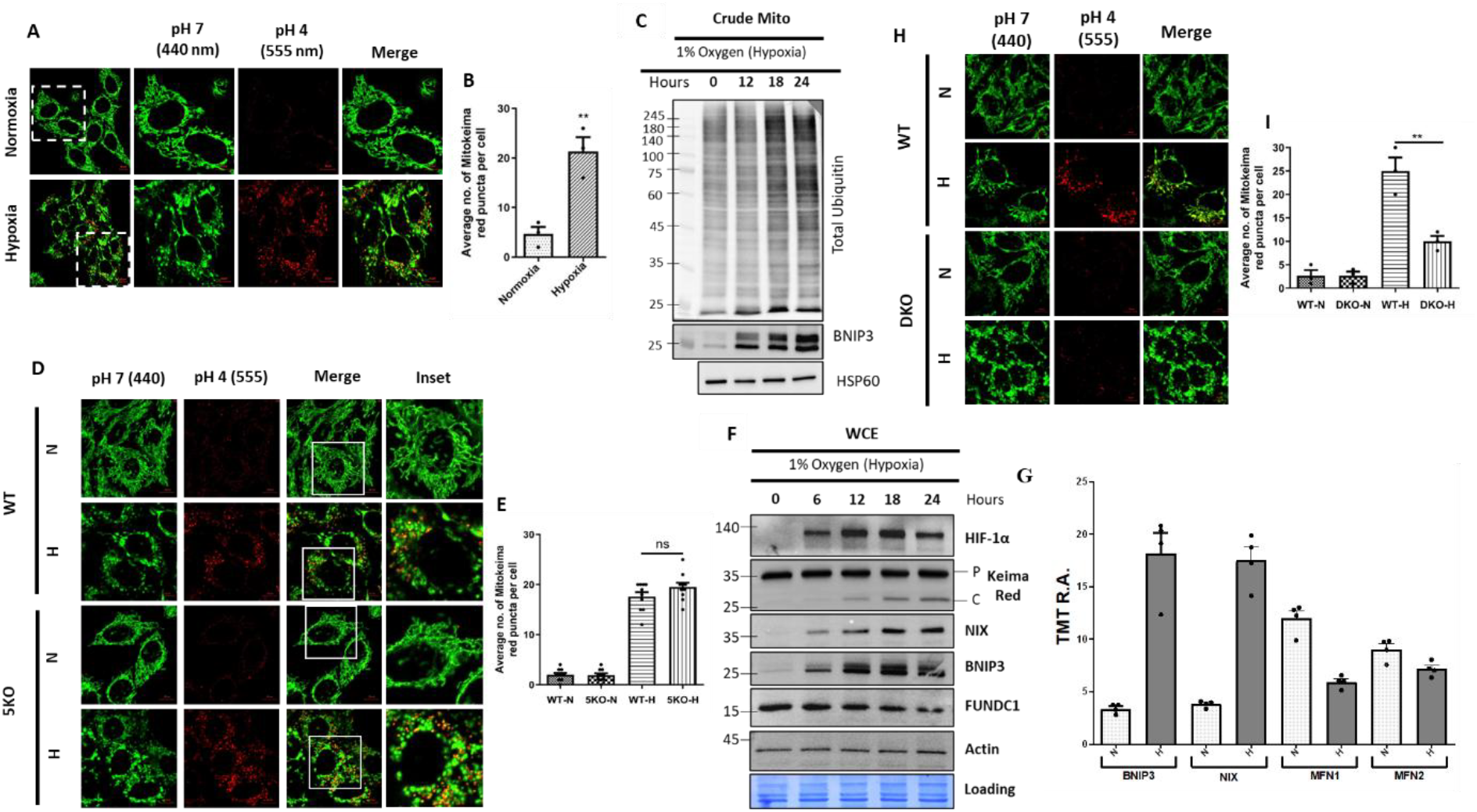
Effects of hypoxia on mitochondria. **A**. Representative images of HeLa cells stably expressing matrix-targeted-Keima-XL (mitoKeima), grown under normoxic or hypoxic conditions. Functional mitophagy was monitored by dual excitation of mitokeima with 440 nm and 555 nm lasers and evaluating the relative shift in mitoKeima excitation. **B**. Quantification of mitophagy by the average number of red puncta per cell in ten independent microscopic fields treated as in panel A. The data is represented as mean±SEM of triplicate experiments. **C**. Immunoblots of total ubiquitin, BNIP3, and HSP60, in crude mitochondria fractions isolated from HeLa cells that were exposed to hypoxia as per the indicated time points. **D**. Representative images of WT and 5KO (deleted for p62/SQSTM1, NDP52, Optineurine, NBR1 and TAX1BP1) cells stably expressing mitoKeima exposed to hypoxia or normoxia. Functional mitophagy was evaluated by the relative shift in mitoKeima excitation. Scale bar: 10 µm. **E**. Quantification of mitophagy by the average number of red puncta per cell in ten independent microscopic fields treated as in panel D. The data is represented as mean±SEM of triplicate experiments. **F**. Immunoblots of clarified whole cell extracts (WCE) from cells stably expressing mitoKeima and exposed to hypoxia for the indicated time points. Mitophagy was estimated by immunoblotting the mitoKeima cleavage using anti-Keima Red antibody: P, Precursor; C, Cleaved. **G**. Crude mitochondria were isolated from HeLa cells following exposure to hypoxic and Normoxic conditions, and proteins were quantified through Tandem mass tag (TMT) Mass Spectrometry. Quantified levels (average from quadruple samples) of relevant proteins are displayed in the bar graph as mean±SEM. **H**. Representative images of WT and DKO (deleted for BNIP3 and NIX) HeLa cells stably expressing mitoKeima, grown under normoxic or hypoxic conditions. Functional mitophagy was evaluated by the mitoKeima assay as mentioned above. Scale bar: 10 µm. **I**. Quantification of mitophagy by the average number of red puncta per cell in ten independent microscopic fields treated as in panel H and the data is represented as mean±SEM of triplicate experiments.

The increased level of mitochondrial ubiquitination prompted us to investigate whether it plays a role in mitochondrial clearance through mitophagy. To test this, we measured mitophagy in a penta knockout (5KO) HeLa cell line lacking five well-known ubiquitin-binding autophagy receptors (p62/SQSTM1, NDP52, Optineurine, NBR1 and TAX1BP1; **EV1D**)^23,24^ stably expressing mitoKeima. After 24 hours of hypoxia exposure, the mitoKeima reporter showed no significant difference in mitophagy levels between the WT and 5KO cells **(Figure 1D, 1E)**. We further evaluated the potential UPS involvement on mitophagy using inhibitors of key enzymes (TAK-243^25^ for the E1 ubiquitin-activating enzyme UBA1, CB-5083^26^ targeting the AAA ATPase p97, MG-132^27^ to inhibit the proteasome, and Bafilomycin A1^28^ for lysosome/autophagy). Only the lysosomal inhibitor Bafilomycin blocked mitophagy **(EV1E)**, supporting the notion that hypoxia-induced mitophagy occurs independent of mitochondrial ubiquitination.

Next, we examined the role of ubiquitin-independent receptors in hypoxia-induced mitophagy. The cellular levels of BNIP3, NIX, and FUNDC1 were monitored over time following the onset of hypoxia exposure. As mitochondria fragmented and were delivered to lysosomes, the resulting cleaved mitoKeima confirmed the progression of mitophagy (**Figure 1F**). Along with the induction of mitophagy, both NIX and BNIP3 levels gradually increased over 24 hours, while FUNDC1 showed a different behavior (**Figure 1F**). Crude mitochondrial fractions were isolated from cells under normoxic and hypoxic conditions. Proteins in these samples were subjected to tryptic digestion and were then differentially labeled using Tandem Mass Tag ^29^. The resulting proteomes, revealed a significant increase in the steady-state levels of BNIP3 and NIX proteins in hypoxic cells; this increase was not due to mitochondrial biogenesis, as outer mitochondrial membrane (OMM) proteins, like MFN1 and MFN2 (Mitofusin) decreased (**Figure 1G**). The loss of mitofusins can also explain the breakdown of the mitochondrial network and the abundance of fragmented mitochondria, which favors mitophagy^30^. To confirm BNIP3/NIX’s role in hypoxia-induced mitophagy, we generated double knockout (DKO) HeLa cells by CRISPR/Cas9 technology **(EV1F)**. A marked decrease in hypoxia-induced mitophagy was observed in the DKO **(Figure 1H, 1I)**, indicating that BNIP3 and NIX are the main mitophagy receptors under these conditions.

### Linear Ubiquitin chains associate with mitochondria in response to hypoxia

The significant buildup of polyubiquitin at mitochondria during hypoxia is puzzling, considering that hypoxia-induced mitophagy seems to be independent of ubiquitin. To explore this response, we mapped the mitochondria-associated ubiquitin landscape following hypoxia exposure. We employed TMT-based comparative analysis on crude mitochondria and enriched for peptides bearing the signature ubiquitin remnants on modified lysines “KεGG” (**Figure 2A**)^31^. The greatest change measured by this approach was for a mixed M1-K6 modified ubiquitin-derived peptide, which increased three-fold **(Figure 2B)**. In contrast, there was no similar increase in unbranched K6 linkages (**Figure 2B**). Since our peptide enrichment method involved capturing KεGG remnants with an antibody, it was limited to detecting isopeptide bonds; thus, we could not directly enrich for linear (head-to-tail) M1 linkages. Nonetheless, mixed chains containing both M1 and K6 linkages could be enriched, because they contain a KεGG motif within the same peptide that also harbors the GGM sequence remnant after tryptic cleavage of M1 linkages **(Figure 2A)**.

**Figure 2:**
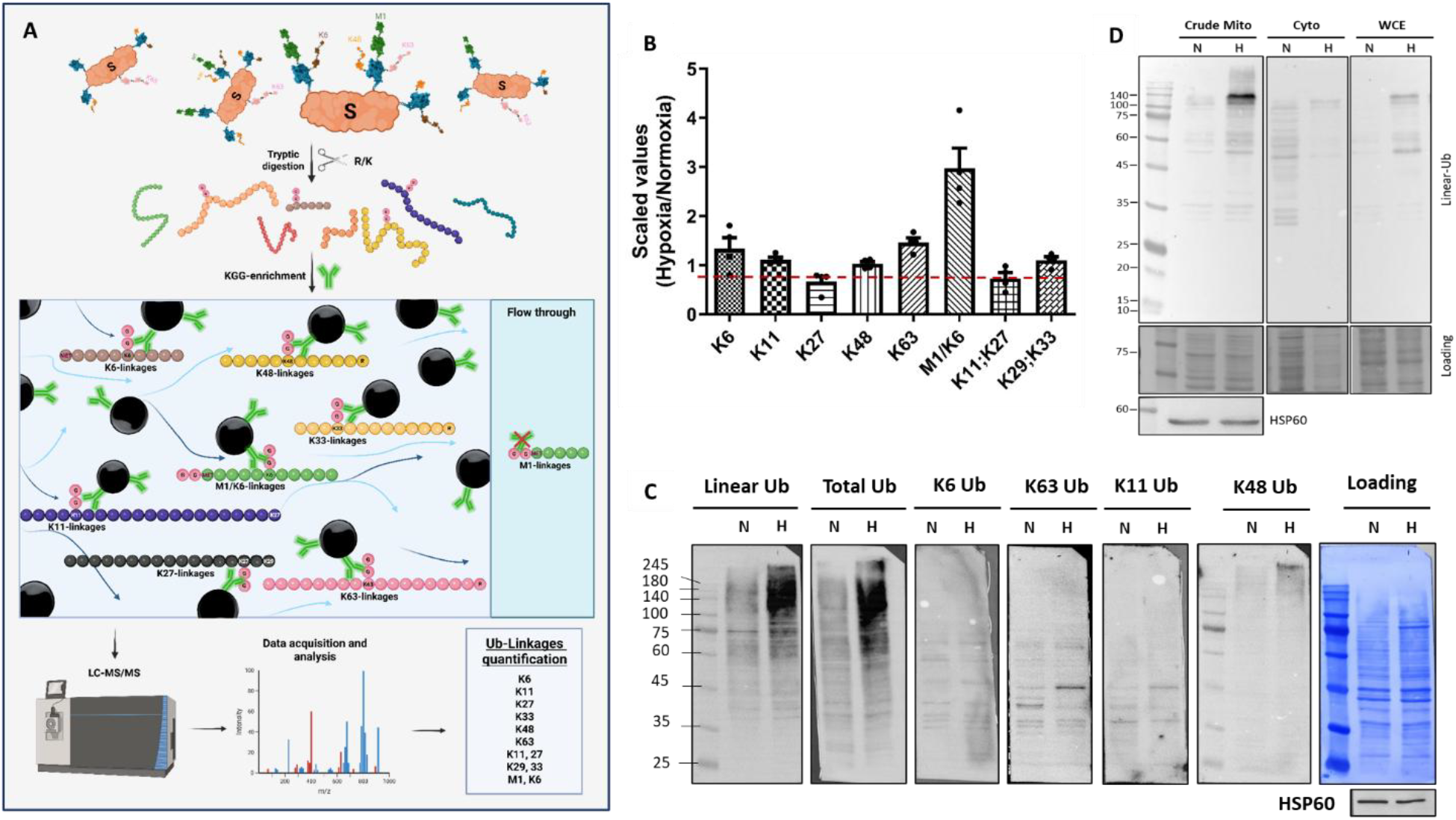
linear Ubiquitin chains associated with mitochondria in response to hypoxia. **A**. A schematic representation of mitochondrial KεGG enrichment. Isolated crude mitochondria are differentially isotopically labelled (Tandem Mass Tagged), and then subject to tryptic digestion. Peptides with residual diglycine modifications on Lysine side chains (KεGG) are selectively enriched with a specific antibody for KεGG. Peptides are analyzed using liquid chromatography-tandem mass spectrometry (LC-MS/MS) to identify different types of isopeptide linkages on ubiquitin-derived peptides. Note that the peptide products of trypsin digestion of M1 linear ubiquitin cannot be detected with KεGG antibody. **B**. Changes in ubiquitin landscape at mitochondria following hypoxia. Crude mitochondria isolated from hypoxic and normoxic cells were differently labeled and analyzed as in Scheme A. Ratios of linkage abundance between hypoxic and normoxic conditions are plotted for each linkage type. K6, K11, K27, K48, and K63 represent ubiquitination on a single lysine of ubiquitin, whereas K11;K27, K29;K33 represent dual modifications on a single peptide. M1/K6 reflects a KεGG at the sixth lysine of the ubiquitin sequence, to which two glycine’s are appended to the N-terminus. **C**. Crude mitochondria were isolated from WT HeLa cells following exposure to hypoxia at 1% for 24 hours. Normoxic (N) and hypoxic (H) crude mitochondrial extracts were immunoblotted with ubiquitin linkage-specific antibodies. HSP60 serves as a loading control for mitochondria. **D**. Clarified whole cell extract (WCE) from cells treated as in C was fractionated into cytosolic and crude mitochondria. The samples were immunoblotted for M1 Linear-Ub. HSP60 serves as a loading control for mitochondria.

Immunoblotting with linkage-specific antibodies provided direct evidence of a robust accumulation of M1 linear chains in crude mitochondrial samples (**Figure 2C**). In this regard, M1 linear ubiquitin chains stood out compared to other tested ubiquitin linkages. When clarified whole cell extract (WCE) was fractionated into cytosolic and crude mitochondrial fractions, the main accumulation of M1 linkages was found in the crude mitochondrial fractions (**Figure 2D**). Overall, these results suggest that linear ubiquitin chains are a major component of hypoxia-induced ubiquitination associated with mitochondria.

### Hypoxia-induced linear ubiquitin chains are triggered by reactive oxygen species

Next, we aimed to identify what triggers the appearance of linear ubiquitin chains at mitochondria in response to hypoxia. During hypoxia, electron transfer becomes less efficient due to the limited availability of molecular oxygen (the terminal electron acceptor of the electron transport chain), leading to the production of reactive oxygen species (ROS), mainly by complex I and III^32^. Using fluorescence microscopy and flow cytometry, we confirmed that hypoxia increases mitochondrial ROS (**Figure 3A,B**). A notable reduction in MitoSOX Red fluorescence was observed upon MQ treatment (a mitochondria-targeted antioxidant shown to preserve mitochondrial function^33^), supporting the role of hypoxia in mitochondria-derived ROS production **(Figure 3B,C)**. In addition to mitochondria-related ROS, the hypoxic response involves several phenomena: HIF-1α stabilization, increased mitochondrial ubiquitin, autophagy receptor accumulation, mitophagy, and elevated antioxidants (**Figure EV3A**,**B**,**C**). As expected, MQ counteracts these responses (**Figure EV3A**,**B**). Collectively, these observations suggest that mitochondrial ROS play a central role in hypoxia-induced mitochondrial ubiquitination and mitophagy.

**Figure 3:**
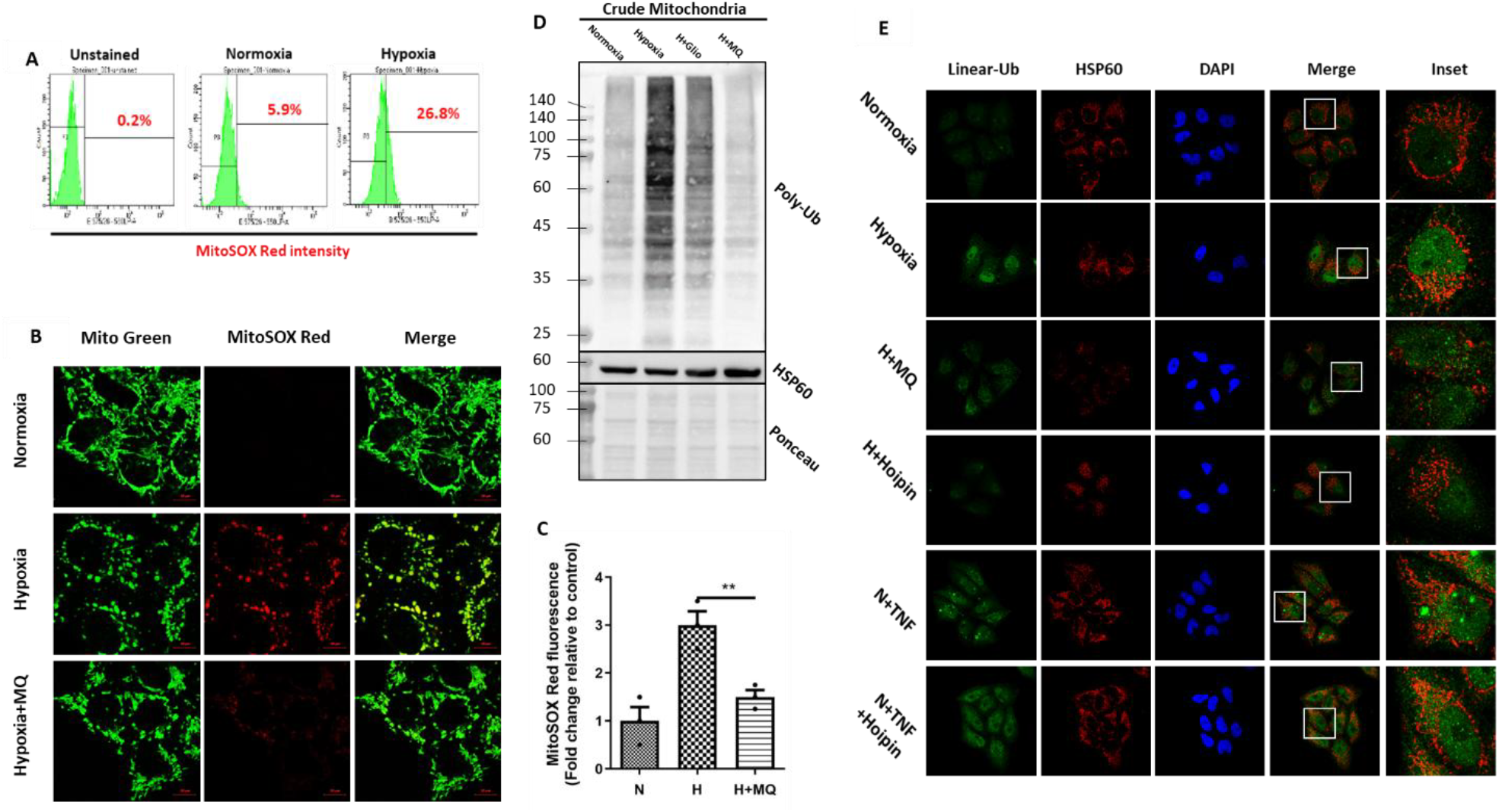
Hypoxia associated linear Ub chains are driven by mitochondrial ROS. **A**. The cells either grown under normoxia or exposed to hypoxia (1% for 24H) were stained with MitoSOX Red dye (detecting superoxide in the mitochondrial context) and analyzed by flow cytometry. Histogram analysis was performed to ascertain the percentage of cell population with MitoSOX Red staining. **B**. HeLa cells were grown under normoxia, hypoxia (1% for 24H) or hypoxia+MQ (1 µM MQ) were stained with MitoTracker Green and MitoSOX Red dyes and imaged by confocal microscope. **C**. Histograms of (B) showing the MitoSOX intensity fold change compare to normoxia. ^**^ p<0.01. **D**. Crude mitochondria were isolated from WT HeLa cells grown in normoxia, hypoxia, hypoxia + MQ (1 µM) or Hypoxia + Gliotoxin (1 µM). The extracts were analyzed by immunoblotting for the detection of total ubiquitin linkages. HSP60 confirms equal amount of loaded mitochondrial fraction. **E**. HeLa cells growing in normoxia, hypoxia, hypoxia + MQ (1 µM), hypoxia + Hoipin-8 (30 µM), normoxia + TNF (20 ng/mL) or normoxia +TNF + Hoipin-8. TNF treatment serves as a positive control for the detection of M1-Ub. All samples were stained with DAPI (nucleus), linear-ubiquitin and mitochondria and imaged by confocal microscope.

Given the pivotal role of mitochondrial ROS in the accumulation of total ubiquitin, we were prompted to ask whether the specific linear ubiquitin buildup is mitoROS-dependent. Analysis of crude mitochondrial fractions treated with Gliotoxin, an inhibitor of LUBAC, showed a decrease in the ubiquitin signal associated with mitochondria in response to hypoxia **(Figure 3D)**. This aligns with a large portion of mitochondria-associated ubiquitin chains of linear linkages. Since Gliotoxin has been documented to cause certain side effects^34^, for further experiments, we used Hoipin-8, a more specific LUBAC inhibitor^35^. To obtain direct evidence that mtoROS triggers linear ubiquitin chain buildup, we performed co-immunofluorescence using a specific antibody for M1-Ub. Both MQ and Hoipin-8 significantly suppressed the accumulation of linear Ub chains in response to hypoxia, further supporting the role of mitochondria-derived ROS in initiating this chain buildup **(Figure 3E)**. So far, our findings indicate that hypoxia-induced mitochondrial ROS activate two separate pathways: mitophagy and linear Ub chain formation. Since we demonstrated that these pathways are independently triggered by ROS, an open question remains: what is the outcome of mitochondria-associated linear Ub chains?

### Hypoxia-driven linear Ubiquitin chains induce canonical NF-κB signaling

Linear ubiquitin chains have several physiological roles, including immune regulation, inflammation, and cell death^36^. Specifically, published literature has shown that linear ubiquitin chains regulate canonical NF-κB signaling, resulting in a proteomic shift driven by NF-κB-induced genes^37,38^. We asked whether these hypoxia-induced linear Ub chains activate NF-κB signaling. We identified a significant reduction in the three NF-κB inhibitor subunits (NFκBIA, NFκBIB, and NFκBIE) associated with crude mitochondria in hypoxic samples **(Figure 4Aa)**. Furthermore, we observed a substantial increase in Interleukin-6 (IL-6) and FOS protein levels, which are products of key NF-κB target genes **(Figure 4Ab)**. These findings suggest that hypoxia activates NF-κB signaling.

**Figure 4:**
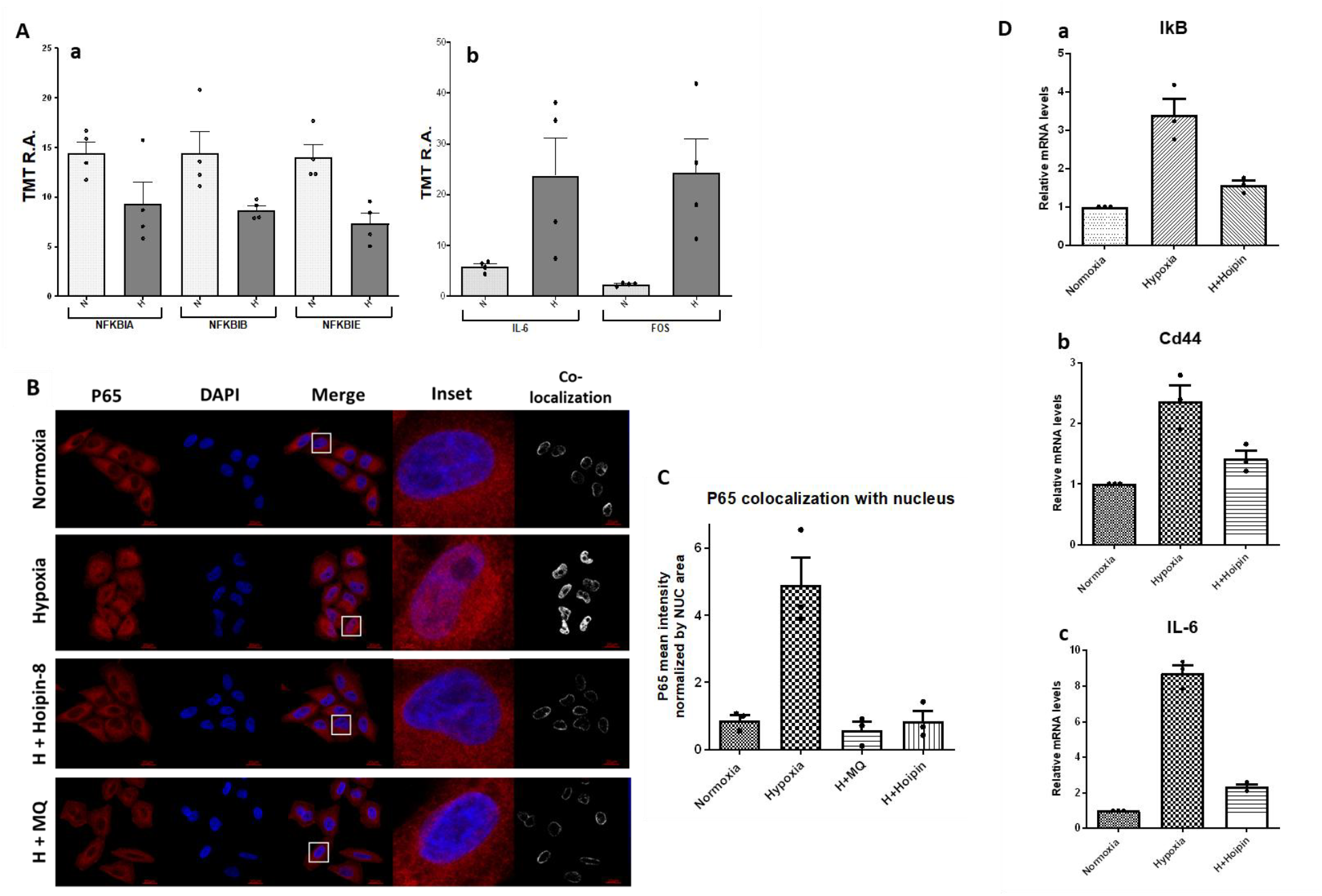
Induction of canonical NF-κB signaling through mitochondria-derived ROS. HeLa cells were exposed to 1% Oxygen (Hypoxia) for either 24 hours or as per the indicated time points. A. Crude mitochondria were isolated from HeLa cells following exposure to hypoxic and Normoxic conditions and then analyzed through Tandem mass tag (TMT) Mass Spectrometry. The relative abundance of the following NF-KB-related proteins is plotted as a bar graph: (a) IkB subunits. (b) cytokines. B. HeLa cells were divided into four sample plates: Normoxic conditions (Control); Hypoxic conditions; Hypoxia + 1uM MitoQuinone; Normoxia + 25ng/mL TNF-alpha and H + 30ng/mL Hoipin-8. All samples were stained with nucleus (Blue) and P65 (Red). C. Histogram of (B) showing the relative intensity of P65 localized to the nucleus, normalized by nucleus area. Individual dots represent biologically independent D. RT-qPCR analysis of NF-κB target genes in HeLa cells under normoxia, hypoxia or hypoxia + HOIPIN-8. mRNA levels of IκBα, CD44, and IL-6 were quantified and normalized to GAPDH. Hypoxia increased NF-κB target gene expression, which was reduced by HOIPIN-8 treatment. Data represent mean ± SEM from three independent experiments.

For NF-κB activation, we expect to see the p65 subunit (RelA) translocate to the nucleus. Indeed, significant nuclear localization of p65 in hypoxic cells was visualized by immunofluorescence **(Figure 4B,C)**. Treatment with MQ disrupted the nuclear localization of p65, indicating a pivotal role of Mito-ROS in activating NF-κB signaling. Additionally, HOIPIN-8 reduced the nuclear presence of p65 **(Figure 4B,C; EV4)**, suggesting that linear chains contribute to hypoxia-induced NF-κB activation. Using Hoipin-8 on hypoxic cells also suppressed the expression of canonical NF-κB target genes, as shown by mRNA levels (**Figure 4D**). These findings outline the sequence of events after hypoxia exposure, leading to the activation of the inflammatory response through mitoROS-driven M1-linear ubiquitin chains.

## Discussion

In this study, we provide new insights into how cells respond to hypoxia, focusing especially on the role of hypoxia-driven mitochondrial ROS (mitoROS) in the buildup of linear ubiquitin chains and activation of NF-κB. mitoROS affects the formation of ubiquitin conjugates, mainly with linear ubiquitin linkages near or on mitochondria, which then activate NF-κB. NF-κB signaling is well known for its involvement in inflammation and stress responses, with linear ubiquitin chains acting as key mediators of this pathway. Here, we observe a prolonged response compared to the typical short-term inflammatory response. Interestingly, our results indicate that linear ubiquitin chains can be formed through an internal trigger – mitoROS – distinct from the usual activation by external signals, such as cytokines, which require outer membrane receptors.

Mitochondrial ROS are an inevitable consequence of hypoxia, which provokes several adaptive mechanisms. One such mechanism is mitophagy, which reduces mitoROS levels by degrading mitochondria when oxygen is scarce. A well-defined pathway for mitophagy involves the promiscuous ubiquitination of mitochondrial outer membrane proteins, which are then recognized by ubiquitin-dependent mitophagy adaptors. Although we observed increased mitophagy and higher ubiquitin levels under hypoxia, we found that hypoxia-induced mitophagy occurs through the ubiquitin-independent receptor-mediated pathway. Thus, mitoROS influence both the extent of mitophagy and the formation of linear chains, but these two pathways seem to operate in parallel, serving complementary roles in adaptation.

While this study offers valuable insights, there are several key areas for future research. First, focusing on the specific localization of linear ubiquitin chain buildup and the associated proteins is crucial to understanding the mechanistic pathway. Although we have identified several important factors involved, the full mechanism behind the ubiquitin accumulation at the mitochondria remains to be clarified. Additionally, although our experiments were performed in cultured HeLa cells, which provide useful information, extending this work to solid tumor models and non-tumorigenic cell lines is essential. Since hypoxia plays a central role in the tumor microenvironment, these models could offer a more comprehensive understanding of the pathway in a physiologically relevant setting. Exploring how hypoxia triggers NF-κB activation may reveal important insights into how cancer cells adapt and survive under low oxygen conditions. This could lead to the identification of new therapeutic targets to fight solid tumor survival. Furthermore, our findings underscore the wide-ranging role of linear ubiquitin chains in regulating various stress response mechanisms.

## Materials and methods

### Cell culture

HeLa cells (wild type, ATG5 knockout, penta knockout, BNIP3/NIX double knockout) were cultured in DMEM containing 10% Fetal bovine serum (FBS), 2 mM Glutamine, 1 mM Sodium Pyruvate and 1% antibiotics (Penicillin and Streptomycin). Wild type, ATG5 knockout, and penta knockout HeLa cells were kindly provided by Prof. Richard Youle (NINDS, NIH, USA). The BNIP3/NIX double knockout cell lines were generated by CRISPR-Cas9 approach. Following targeting sequences were used to design the guide RNAs: **GGAGAGAAAAACAGCTCAC**; BNIP3: NIX: **CAGGACAGAGTAGTTCCAG**. The complementary guide RNA oligonucleotides were annealed together to generate a guide RNA duplex, which is ligated to BpiI digested pSp-Cas9-BB-2A-Puro (PX459) V 2.0 vector (a gift from Feng Zhang; Addgene Plasmid #62988). The plasmids were confirmed by restriction digestion and sequence verified. Wild type HeLa cells were transfected with the CRISPR plasmids using the X-treme Gene HP transfection reagent (Roche), followed by selection with Puromycin (2 µg/ml) for two consecutive cycles. The BNIP3/NIX double knockout cell line was generated by co-transfection with equimolar amounts of BNIP3 and NIX CRISPR plasmids. Following puromycin selection, the individual clones were expanded, screened by immunoblotting and target gene microdeletion by sequencing.

### Mitophagy assessment

The extent of mitophagy was detected either by mitoKeima construct (by fluorescence microscopy) or by mtx-mKeimaXL probe (by immunoblotting). mitoKeima (mitochondria-targeted Keima) is a ratiometric, pH-sensitive fluorescent protein resistant to lysosomal proteases which allows rapid and faithful determination of mitophagy^22^. pCHAC-mitoKeima retroviral construct was kindly gifted by Dr. Michael Lassarou (Monash University, Australia; Addgene plasmid # 72342). mitoKeima expressing stable lines were generated by retroviral transduction. mitoKeima expressing cells were cultured in 35 mm confocal (live-cell imaging) plates with a glass bottom. At the end of the desired treatment period, the media was replaced with warm live-cell imaging solution (Invitrogen), containing 25 mM Glucose and 10% FBS. The cells were imaged under a Zeiss LSM 710 fluorescence confocal microscope at 63X oil immersion objective while maintaining in an incubation chamber at 37^°^C and 5% CO_2_. For each sample, 10 distinct microscopic fields were captured, each containing at least 5-10 cells. The extent of mitophagy was indicated as the average number of red-only mitoKeima puncta per cell.

Alternatively, the extent of mitophagy was determined by the mtx-mKeima^XL^ probe, which was kindly provided by Dr. Alban Ordureau (Memorial Sloan Kettering Cancer Center, New York, USA). mtx-mKeima^XL^ expressing stable cell lines were either generated by lentiviral transduction or the cells were transiently transfected. The extent of mitophagy was determined semi-quantitatively by detecting the cleaved Keima protein by immunoblotting and calculating the ratio of cleaved to uncleaved mtx-mKeima^XL^.

### Isolation of crude mitochondria

Cells were washed twice with cold PBS and then scraped in ice-cold HM buffer (250 mM Sucrose, 1 mM EDTA, 1 mM EGTA, 20 mM HEPES pH 7.4) containing 100 mM 2-Chloracetamide and protease inhibitor cocktail. The cell suspension was centrifuged at 500g for 5 minutes to pellet down the cells, followed by resuspension in fresh HM buffer containing 100 mM 2-Chloracetamide and protease inhibitor cocktail. The cells were homogenized in a precooled Dounce homogenizer (Teflon in glass type) with ∼40 strokes on ice and the resulting homogenate was centrifuged twice at 1400g for 10 minutes at 4^°^C in tandem. The resulting supernatant was further centrifuged at 10,000g for 10 minutes at 4^°^C. The crude mitochondrial pellet thus obtained was washed at least 3 times with ice-cold HM buffer.

### Protein extraction and western blotting

The cells were washed twice with cold PBS and scraped directly into ice-cold lysis buffer (50 mM Tris, pH 7.4, 150 mM NaCl, 0.5% NP-40) containing protease inhibitor cocktail. The cell lysates were pipetted and vortexed vigorously, stored on ice briefly followed by centrifugation at 13,000g for 15 minutes at 4^°^C. The clear supernatant was transferred to fresh tube followed by protein estimation by Bradford assay. Equal amounts of cell lysates from each sample were resolved by SDS-PAGE followed by electroblotting onto PVDF membranes. The blots were blocked in skimmed milk and then incubated with primary antibodies either for 1 hour at room temperature or overnight at 4^°^C on a shaker followed by incubation with respective secondary antibodies. The signals were detected using chemiluminescence detection system (Fusion Pulse).

Following antibodies were used: HIF-1α (ab179483), LC-3B (ab51520) and antibodies were obtained from Abcam. Cleaved caspase-3 (Cat#9664) and HSP60 Rb (Cat#12165) antibodies were purchased from Cell Signaling. LAMP-1 (sc-20011), VDAC1 (sc-8828), TOM20 (sc-17764), MFN-1 (sc-166644), MFN-2 (sc-100560), DRP1 (sc-271583), BNIP3 (sc-56167), NIX (sc-166332), Ubiquitin (sc-8017) antibodies were bought from Santacruz Biotechnology. FUNDC1 (PA5-58535) antibody was purchased from Thermo. Keima Red antibody (M182-3M) was obtained from MBL. Following antibodies were obtained from Milipore: Ubiquitin Lysine48-linkage specific (Cat#05-1307), Ubiquitin Lysine63-linkage specific (Cat#1308) and Phospho-Ubiquitin specific (Ser65) (Cat#ABS 1513-I), P65 (RelA) (SC-8008), M1-Ub linkages specific for WB (LUB9; Merck), M1-Ub linkages specific for IF (1E3; zoomab), HSP60 Ms (Proteintech; #66041).

### Evaluation of mitochondrial ROS levels

Mitochondrial ROS was detected using the mitochondrial superoxide indicator dye MitoSOX Red as described earlier^33^. Briefly, cells were growing in confocal plates were stained with 200 nM MitoTracker Green FM and 5 µM MitoSOX Red (Invitrogen) for 15 minutes at 37^°^C in warm serum-free DMEM. The staining solution was replaced with Live-cell imaging solution after washing the cells twice with warm PBS. The images were acquired on a Zeiss 710 confocal microscope while maintaining the cells in the incubation chamber. At least 100 cells across 10 random microscopic fields were analyzed for quantification of MitoSOX Red fluorescence intensity. The data analysis was performed using the ImageJ software (NIH) and indicated as fold change of MitoSOX Red fluorescence intensity relative to control.

### FACS-based evaluation of mitochondrial ROS

To quantify the mitochondrial ROS accumulation following hypoxia, the cells were stained with 5 μM MitoSOX Red for 10 minutes at 37°C in dark followed by acquisition on FL-2 channel of BR LSRII flow cytometer (Becton Dickinson, USA) and data was analyzed on Cell Quest software (BD Biosciences).

### Immunofluorescence

Cells grown on coverslips were gently washed with warm PBS twice and then fixed using warm 4% paraformaldehyde. Post fixation, the cells were permeabilized with 1% Triton X-100 for 5 minutes at room temperature and then blocked in 5% BSA for 1 hour. The cells were then exposed to the desired primary antibodies either for 1 hour at room temperature or for overnight at 4^°^C. Following this, the coverslips were washed thrice with PBST and then cells were incubated with the respective fluorophore-conjugated secondary antibodies for 1 hour at room temperature in dark. The coverslips were washed again with PBST thrice and then mounted with Fluoromount G (eBiosciences) on glass slides and stored at -20^°^C until image acquisition. The colocalization analysis was performed by using the Profile analysis tool in the ZEN lite (V 3.0) software (Zeiss) by fluorescence intensity line measurement approach.

### rtPCR

Total RNA was extracted from HeLa cells using TRIzol reagent (T9424; Sigma-Aldrich) according to the manufacturer’s protocol. The RNA was subsequently purified and concentrated using the RNA Clean & Concentrator (RCC) kit (R1019; Zymo Research) to ensure removal of contaminants and preservation of RNA integrity. Complementary DNA (cDNA) was synthesized from 400 ng of total RNA using the qScript cDNA Synthesis Kit (95047-025; Quantabio) according to the manufacturer’s instructions. Quantitative real-time PCR (RT-qPCR) was performed with the SYGreen Mix Low ROX kit (PB20.11-20; qPCRBIO) on a QuantStudio 1 Real-Time PCR System (Applied Biosystems) using gene-specific primers (Table X). Thermal cycling conditions were as follows: polymerase activation at 95 °C for 2 min, followed by 40 cycles of denaturation at 95 °C for 5 s and annealing/extension at 60–65 °C for 20–30 s, with fluorescence data acquisition on the FAM channel during annealing/extension. A melt curve analysis was performed at the end of the amplification protocol according to the instrument’s instructions. Relative transcript levels were determined using the ΔΔCt method, normalized to GAPDH.

### Proteomics

#### General Sample Preparation

Protein extracts were subjected to disulfide bond reduction with 5 mM TCEP (room temperature, 10 min) and alkylation with 25 mM chloroacetamide (room temperature, 20 min). To prepare for protease digestion, the methanol-chloroform precipitation method was used. Each sample was mixed with four parts of neat methanol, followed by one part of chloroform, and then three parts of water. The mixture was vortexed after each addition. The sample was centrifuged at 6 000 rpm for 2 min at room temperature and subsequently washed twice with 100% methanol. Samples were resuspended in 100 mM EPPS pH8.5 containing 0.1% RapiGest and digested at 37°C for 1h with LysC protease at a 200:1 protein-to-protease ratio. Trypsin was added at a 100:1 protein-to-protease ratio, and the reaction was incubated for 6 h at 37°C. Samples were acidified with 1% Formic Acid for 15 min and subjected to C18 solid-phase extraction (SPE) (Sep-Pak, Waters).

#### Total proteomics analysis using TMTpro

Tandem mass tag labeling of each sample (100 μg peptide input) was performed by adding 10 μL of the 20 ng/μL stock of TMTpro reagent along with acetonitrile to achieve a final acetonitrile concentration of approximately 30% (v/v). Following incubation at room temperature for 1 h, the reaction was quenched with hydroxylamine to a final concentration of 0.5% (v/v) for 15 min. The TMTpro-labeled samples were pooled together at a 1:1 ratio. The sample was vacuum centrifuged to near dryness and subjected to C18 solid-phase extraction (SPE) (Sep-Pak, Waters).

Dried TMT-labeled sample was resuspended in 100 μL of 10 mM NH4HCO3 pH 8.0 and fractionated using BPRP HPLC^39^. Briefly, samples were offline fractionated over a 90 min run, into 96 fractions by high pH reverse-phase HPLC (Agilent LC1260) through an aeris peptide xb-c18 column (Phenomenex; 250 mm x 4.6 mm) with mobile phase A containing 5% acetonitrile and 10 mM NH4HCO3 in LC-MS grade H2O, and mobile phase B containing 90% acetonitrile and 10 mM NH4HCO3 in LC-MS grade H2O (both pH 8.0). The 96 resulting fractions were then pooled non-continuously into 24 fractions (as outlined in Figure S5 of (Paulo et al., 2016a)^40^) and used for subsequent mass spectrometry analysis. Fractions were vacuum centrifuged to near dryness. Each consolidated fraction was desalted via StageTip, dried again via vacuum centrifugation, and reconstituted in 5% acetonitrile, 1% formic acid for LC-MS/MS processing.

Mass spectrometry data were collected using an Orbitrap Eclipse Tribrid mass spectrometer (Thermo Fisher Scientific, San Jose, CA) coupled to an UltiMate 3000 RSLCnano system liquid chromatography (LC) pump (Thermo Fisher Scientific) and the FAIMS Pro Interface. Peptides were separated on a 100 μm inner diameter microcapillary column packed in house with ∼30 cm of HALO Peptide ES-C18 resin (2.7 µm, 160 Å, Advanced Materials Technology, Wilmington, DE) with a gradient consisting of 5%–24% (0-60 min), 24%–38% (60-85min) (ACN, 0.1% FA) over a total 95 min run at ∼500 nL/min. For analysis, we loaded 1/10 of each fraction onto the column. Proteome analysis used Multi-Notch MS^3^-based TMT quantification^41^, combined with a newly implemented Real-Time Search analysis software^42^, and the FAIMS Pro Interface (using previously optimized 3 CV parameters^43^), to reduce ion interference. The scan sequence began with an MS^1^ spectrum (Orbitrap analysis; resolution 120,000 at 200 Th; mass range 400−1500 m/z; automatic gain control (AGC) target 4×10^5^; maximum injection time 50 ms). Precursors for MS^2^ analysis were selected using a cycle type of 1.25 sec/CV method (FAIMS CV=-40/-60/-80). MS^2^ analysis consisted of collision-induced dissociation (quadrupole ion trap analysis; Rapid scan rate; AGC 1.0×10^4^; isolation window 0.5 Th; normalized collision energy (NCE) 35; maximum injection time 35 ms). Monoisotopic peak assignment was used, and previously interrogated precursors were excluded using a dynamic window (180 s ±10 ppm). Following acquiring each MS^2^ spectrum, a synchronous-precursor-selection (SPS) API-MS^3^ scan was collected on the top 10 most intense ions b or y-ions matched by the online search algorithm in the associated MS^2^ spectrum^42^,^44^. MS^3^ precursors were fragmented by high energy collision-induced dissociation (HCD) and analyzed using the Orbitrap (NCE 45; AGC 2.5×10^5^; maximum injection time 200 ms, resolution was 50,000 at 200 Th). The closeout was set at two peptides per protein per fraction, so MS^3^s were no longer collected for proteins with two peptide-spectrum matches (PSMs) that passed quality filters^45^.

#### Immunoprecipitation of diGLY-Containing Peptides

diGLY capture was performed largely as described^46^. The diGly monoclonal antibody (Cell Signaling Technology; D4A7 clone) (32 μg antibody/1 mg peptide) was coupled to Protein A Plus Ultralink resin (1:1 μL slurry/ μg antibody) (Thermo Fisher Scientific) overnight at 4°C before its chemical cross-linking reaction. Dried peptides (1 mg starting material) were resuspended in 1.5 mL of ice-cold IAP buffer [50 mM MOPS (pH 7.2), 10 mM sodium phosphate and 50 mM NaCl] and centrifuged at maximum speed for 5 min at 4°C to remove any insoluble material. Supernatants (pH ∼7.2) were incubated with the antibody beads for 2 hr at 4°C with gentle end-over-end rotation. After centrifugation at 215 × g for 2 min, beads were washed thrice with ice-cold IAP buffer and twice with ice-cold PBS. The diGLY peptides were eluted twice with 0.15% TFA, desalted using homemade StageTips and dried via vacuum centrifugation, before TMTpro labeling.

#### diGLY proteomics analysis using TMTpro

TMTpro-labeled diGLY peptides were fractionated according to the manufacturer’s instructions using High pH reversed-phase peptide fractionation kit (Thermo Fisher Scientific) for a final 6 fractions and subjected to C18 StageTip desalting before MS analysis.

Mass spectrometry data were collected using an Orbitrap Eclipse Tribrid mass spectrometer (Thermo Fisher Scientific, San Jose, CA) coupled to an UltiMate 3000 RSLCnano system liquid chromatography (LC) pump (Thermo Fisher Scientific). Peptides were separated on a 100 μm inner diameter microcapillary column packed in house with ∼30 cm of HALO Peptide ES-C18 resin (2.7 µm, 160 Å, Advanced Materials Technology, Wilmington, DE) with a gradient consisting of 3%–20% (0-100 min), 20%–35% (100-140min) (ACN, 0.1% FA) over a total 155 min run at ∼500 nL/min. For analysis, we loaded 1/2 of each fraction onto the column. For analysis, we loaded half of each fraction onto the column. The scan sequence began with an MS^1^ spectrum (Orbitrap analysis; resolution 120,000 at 200 Th; mass range 400−1300 m/z; automatic gain control (AGC) target 7.5×10^5^; maximum injection time 100 ms). Precursors for MS^2^ analysis were selected using a cycle type of 1.25 sec/CV method (FAIMS CV=-40/-60/-80). MS2 analysis consisted of high energy collision-induced dissociation (HCD) (Orbitrap analysis; resolution 50,000 at 200 Th; isolation window 0.7 Th; normalized collision energy (NCE) 38; AGC 2×10^5^; maximum injection time 300 ms). Monoisotopic peak assignment was used, and previously interrogated precursors were excluded using a dynamic window (120 s ±10 ppm).

#### Data analysis

Mass spectra were processed using a Comet-based (2020.01 rev. 4) in-house software pipeline^47,48^. Spectra were converted to mzXML using a modified version of ReAdW.exe. Database searching included all entries from the Mouse Reference Proteome (2020-03) UniProt database, as well as an in-house curated list of contaminants. This database was concatenated with one composed of all protein sequences in the reversed order. Searches were performed using a 50-ppm precursor ion tolerance for total protein level analysis. The product ion tolerance was set to 0.9 Da (0.03 Da for diGLY searches). These wide mass tolerance windows were chosen to maximize sensitivity in conjunction with Comet searches and linear discriminant analysis^48,49^. TMTpro tags on lysine residues and peptide N termini (+304.207 Da) and carbamidomethylation of cysteine residues (+57.021 Da) were set as static modifications, while oxidation of methionine residues (+15.995 Da) was set as a variable modification. For diGLY dataset search, GlyGly modification (+114.0429 Da) was also set as a variable modification. Peptide-spectrum matches (PSMs) were adjusted to a 1% false discovery rate (FDR)^50^. PSM filtering was performed using a linear discriminant analysis, as described previously^48^, while considering the following parameters: XCorr (or Comet Log Expect), ΔCn (or Diff Seq. Delta Log Expect), missed cleavages, peptide length, charge state, and precursor mass accuracy. For TMTpro-based reporter ion quantitation, we extracted the summed signal-to-noise (S:N) ratio for each TMT channel and found the closest matching centroid to the expected mass of the TMT reporter ion (integration tolerance of 0.003 Da). For protein-level comparisons, PSMs were identified, quantified, and collapsed to a 1% peptide false discovery rate (FDR) and then collapsed further to a final protein-level FDR of 1%. Moreover, protein assembly was guided by principles of parsimony to produce the smallest set of proteins necessary to account for all observed peptides. Ubiquitylation site localization was determined using the AScore algorithm^49^. AScore is a probability-based approach for high-throughput protein phosphorylation site localization. Specifically, a threshold of 13 corresponded to 95% confidence in site localization. Proteins and ubiquitylated peptides were quantified by summing reporter ion counts across all matching PSMs using in-house software, as described previously^48^. PSMs with poor quality, MS^3^ spectra with isolation specificity less than 0.8, or with TMT reporter summed signal-to-noise ratio that was less than 160, or had no MS^3^ spectra were excluded from quantification^51^.

Protein or peptide quantification values were exported for further analysis in Microsoft Excel, GraphPad Prism and Perseus^52^. For whole proteome analysis, each reporter ion channel was summed across all quantified proteins and normalized assuming equal protein loading of all samples. For diGly samples, the data was normalized to each individual protein abundance measured in parallel when available to correct for variation in protein abundance between treatments. Supplemental data Tables list all quantified proteins as well as the associated TMT reporter ratio to control channels used for quantitative analysis.

Annotations for bona fide organellar protein markers were assembled using the proteins which had scored with confidence “very high” or “high” from the HeLa dataset previously published Itzhak D.N.^53^. The following database containing mitochondrial proteins was used: MitoCarta 3.0^54^.

## Author contributions

TK and PS designed and executed the experiments.

AO analysed MS and proteomic data.

JR, NR, and IM aided the experimental procedure and contributed to the interpretation.

TK, PS, and MHG raised the hypothesis, oversaw the progress, and wrote the first draft of the paper.

MHG and AO secured funding.

All authors read and approved the manuscript.

## Acknowledgements and Funding

This work was partially funded by ISF BRG grant 2640/23, and by the European Union ERC grant (UbWan, 101142726) to MHG. This work was funded in part by NIH/NIGMS grant R35 GM156454 (AO) and by the Memorial Sloan Kettering Cancer Center Support Grant P30CA008748 (AO). This manuscript is the result of funding in whole or in part by the National Institutes of Health (NIH). It is subject to the NIH Public Access Policy. Through acceptance of this federal funding, NIH has been given a right to make this manuscript publicly available in PubMed Central upon the Official Date of Publication, as defined by NIH.

Views and opinions expressed are however those of the author(s) only and do not necessarily reflect those of the European Union, the European Research Council Executive Agency or the U.S. National Institutes of Health. Neither the European Union nor the granting authority can be held responsible for them.

## Data Sharing

All relevant dasta is included with the submission. The mass spectrometry proteomics data will be deposited to the open repository MassIVE with the Project accession number (Will be added after submission).

Expanded view data and supplementary information are appended to this paper

