## Supplemental figures for "Hypoxia-induced mitoROS triggers M1 linear ubiquitin chains and activates NF-κB signaling"

**Title: Hypoxia-generated mitoROS induce M1 Linear Ubiquitin Chains and activate NF-κB signaling**

*** These authors contributed equally**

**1 Faculty of Biology, Technion-IIT, Haifa 32000, Israel**

**2 Cell Biology Program, Sloan Kettering Institute, Memorial Sloan Kettering Cancer Center, New York, NY 10065, USA**

**This file contains 3 extended view figures:**

**EV1**

**EV3**

**EV4**

### EV1 for Fig 1

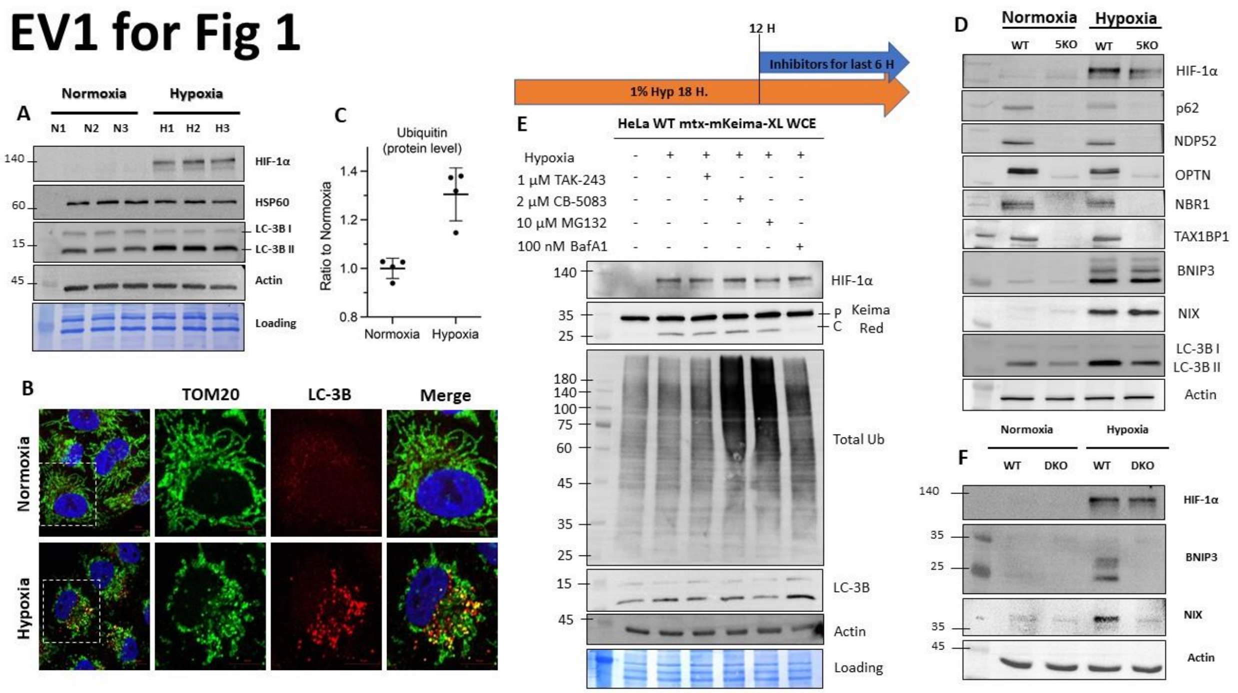


**A.** Immunoblots of whole cell extracts (WCE) from HeLa cells grown under normoxic or hypoxic conditions. Autophagy was estimated by conversion of LC-3BI to LC-3BII. The blot shows a biological triplicate of each condition. **B.** Representative images of HeLa cells stably expressing matrix-targeted-Keima-XL (mitoKeima), grown under normoxic or hypoxic conditions. Functional mitophagy was evaluated by the shift in mitokeima fluorescence. Scale bar: 10 µm. **C.** Quantification of Ubiquitin levels from the proteomics data represented in the form of TMT signal relative to normoxia. (n=4). **D.** Immunoblots of whole cell extracts (WCE) from WT and 5KO HeLa cells grown under normoxic or hypoxic conditions. Autophagy was estimated by conversion of LC-3BI to LC-3BII. The blot shows a biological triplicate of each condition. **E.** Immunoblots of whole cell extracts (WCE) from HeLa cells grown treated as indicated. Mitophagy is estimated by cleavage of Keima-red in each condition. **F**. Immunoblots of whole cell extracts (WCE) from WT or DKO HeLa cells grown under normoxic or hypoxic conditions. The blot shows a biological triplicate of each condition.

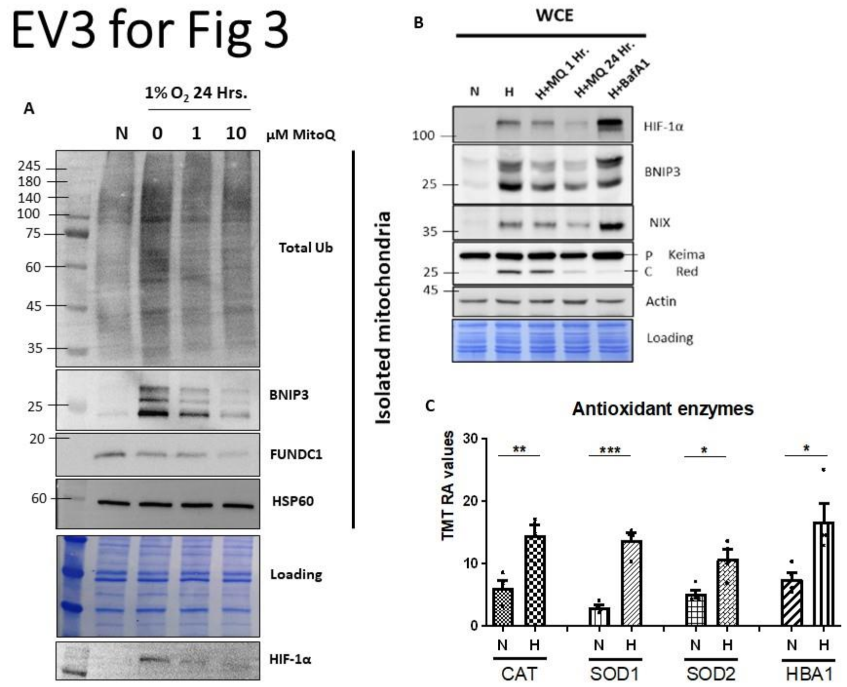


**A.** Immunoblots of crude mitochondria and whole cell extracts from HeLa cells grown under normoxic or hypoxic conditions with different MitoQ concentrations (0uM, 1uM, 10uM). HIF-1a, autophagy receptors, and conjugated ubiquitin were detected by specific antibodies. HSP60 was used as a loading control for mitochondria. **B.** Immunoblots of clarified whole cell extracts (WCE) from HeLa cells expressing mKeima-Red grown under normoxic or hypoxic conditions, treated with MitoQ for the noted times (1Hr, 24Hr) or BAFA1 to arrest global autophagy. HIF-1a and autophagy receptors were measured by specific antibodies. Keima-Red cleavage was used to detect the occurrence of the mitophagy process in all samples. Actin was used as a loading control. **C.** Quadruplicate crude mitochondria were isolated from HeLa cells following exposure to hypoxic or Normoxic conditions and then analyzed through Tandem mass tag (TMT) Mass Spectrometry. The relative levels of notable antioxidants are plotted in a bar graph.

### EV4 for Fig4

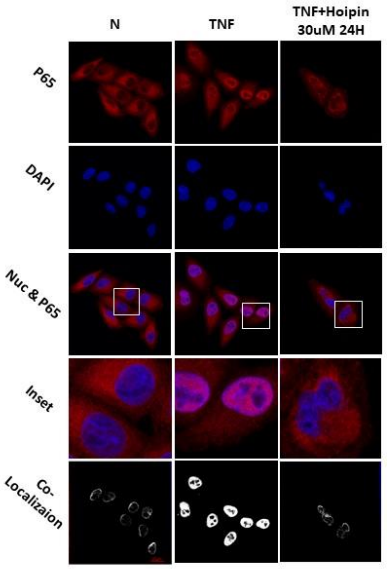


**A.** induction of canonical NF-kB signaling through TNF. HeLa cells were treated with 25ng/ml TNFa for 15 minutes or with 25ng/mL NF-alpha togther with 30ng/mL Hoipin-8. Cells were stained with nucleus (Blue) and P65 (Red).
